# Population Specific Adaptations in Venom Production to Abiotic Stressors in a Widely Distributed Cnidarian

**DOI:** 10.1101/2020.02.28.969204

**Authors:** Maria Y. Sachkova, Jason Macrander, Joachim M. Surm, Reuven Aharoni, Shelcie S. Menard-Harvey, Amy Klock, Whitney B. Leach, Adam M. Reitzel, Yehu Moran

**Affiliations:** Department of Ecology, Evolution and Behavior, Alexander Silberman Institute of Life Sciences, Faculty of Science, The Hebrew University of Jerusalem, Israel; Sars International Centre for Marine Molecular Biology, University of Bergen, Bergen, Norway; Department of Biological Sciences, University of North Carolina at Charlotte, Charlotte, NC, USA; Florida Southern College, Lakeland, FL, USA

## Abstract

*Nematostella vectensis* is a sea anemone (Actiniaria, Cnidaria) inhabiting estuaries over a broad geographic range where environmental conditions such as temperatures and salinity vary widely. In cnidarians, antagonistic interactions with predators and prey are mediated by their venom, which may be metabolically expensive. In this study, we challenged *Nematostella* polyps with heat, salinity, UV light stressors and a combination of all three to determine how abiotic stressors impact toxin expression for individuals collected across this species’ range. Transcriptomics and proteomics revealed that the highly abundant toxin Nv1 was the most downregulated gene under heat stress conditions in multiple populations. Physiological measurements demonstrated that venom is metabolically costly to produce suggesting that downregulating venom expression under stressful conditions may be advantageous. Strikingly, under a range of abiotic stressors, individuals from different geographic locations along this latitudinal cline modulate venom production levels differently in a pattern reflecting local adaptation.

## Introduction

*Nematostella vectensis* is a burrowing sea anemone which specializes in estuarine environments with a unique role as an infaunal predator [1]. These brackish habitats are characterized by dynamic daily and seasonal abiotic conditions, particularly temperature, salinity, and ultraviolet (UV) light [2]. *Nematostella* has a broad geographic range along the Atlantic and Pacific coasts of the United States and southern Canada where the extent of variation in environmental conditions changes by latitude or location within the estuary [1]. *Nematostella* tolerates salinities from ∼8.96 – 51.54‰ and temperatures from −1.5 – 28.5 °C [1, 3] in the field, and laboratory experiments have shown it can acclimate to even broader ranges. Like many coastal invertebrates, *Nematostella* exhibits extensive genetic diversity with significant population genetic structure throughout its range [4, 5]. Current evidence of genetic structure and a life history that likely reduces dispersal between locations (collective egg masses, demersal larvae) is consistent with limited gene flow between estuaries. The combination of geographically structured genetic variation and differences in environmental conditions is the context where we may expect that populations might be adapted to different ranges of environmental parameters [4].

Previous research with *Nematostella* adults from different geographic locations has shown evidence consistent with local adaptation for particular phenotypes or genetic loci [6]. *Nematostella* from locations along the Atlantic coast have temperature dependent growth rates and thermotolerance consistent with a thermal gradient from low to high latitudes [4]. Similarly, different genotypes vary in their tolerance to oxidative stress, which may be related to genetic variation in the transcription factor NF-κB [7] or superoxide dismutase [8]. Although whole genome comparisons have not yet been completed to identify additional loci where genetic variation is structured between populations, a survey of allelic variation for a subset of gene coding loci suggests it may be pervasive [9]. The diversity of genes potentially involved in adaptation to abiotic variation (such as heat, salinity and oxidative stresses) could be large. For example, *Nematostella* has large numbers of heat shock proteins (HSPs) and antioxidant genes [10, 11] likely to be instrumental in mounting a cellular stress response

Species also may adapt to their biotic environment based on the distribution of their prey and predators. In *Nematostella*, as in most cnidarians, antagonistic interactions with predators and prey are largely mediated by the venom system. Adult venom includes multiple toxins produced by nematocytes and ectodermal gland cells [12, 13]. Venom production is regulated throughout the life cycle and correlates with the type of interactions *Nematostella* is exposed to: at the larval non-feeding stage it produces toxins to deter fish predators while as an adult it combines both defensive and offensive toxins [12, 14]. The Nv1 toxin is the major component of the adult venom and is produced by ectodermal gland cells at very high levels as it is encoded by multiple gene copies [15, 16]. Nv1 is lethal even at low doses for crustaceans, which could be either predators or prey [12]. Under controlled diel light conditions, expression of Nv1 is significantly higher during the day when compared with night [17]. In fact, Nv1 is among the most differentially expressed genes in *Nematostella*. The higher expression during the day correlates with an antipredator response to visual predators like shrimp and fish.

Venom production is reported to be metabolically “expensive” [18]. However, the metabolic cost of venom biosynthesis has only been studied in scorpions and snakes [19-21], with no studies in cnidarians. Moreover, even in snakes there are contrasting views regarding the cost of venom to individuals [22]. Regulation of the venom production across *Nematostella*’s life cycle supports a hypothesis that tuning venom composition based on ecological requirements may be advantageous [12]. Otherwise, anemones would be expected to maintain high venom expression regardless of stage or environment. Stress response mechanisms require additional metabolic cost as they involve production of high levels of specialized proteins, such as chaperones [23], depending on the severity of the physiological stress imposed by the environment.

In this study, we hypothesized that there might be an advantage to downregulate venom biosynthesis depending on physiological status of the animal and its exposure to environmental stressors in order to prioritize metabolic resources for supporting the stress response. To test this hypothesis we combined analyses of physiology, gene expression, and proteomics of *Nematostella* adults challenged with combinations of environmental stressors to determine the response in the production of venom. Further, we also compared responses between individuals collected from different locations along this species’ native range (i.e., Atlantic coast of North America) to determine to what extent tradeoffs in stress response and venom production differ between these geographically and genetically isolated populations.

## Results

### Venom production is metabolically expensive

To study whether venom production is metabolically costly, we depleted *Nematostella* venom reserves by mechanical stimulation and measured time-dependent changes in respiration rate, which correlates with metabolic rate. A similar method has been used to assess metabolic cost of venom production in snakes and scorpions [19, 20]. Oxygen intake was measured every hour for five hours following the stimulation (**Fig 1A**). After two hours, oxygen consumption increased by more than 34% and it persisted for the next two hours with a slight decrease by the fifth hour.

**Figure 1.**
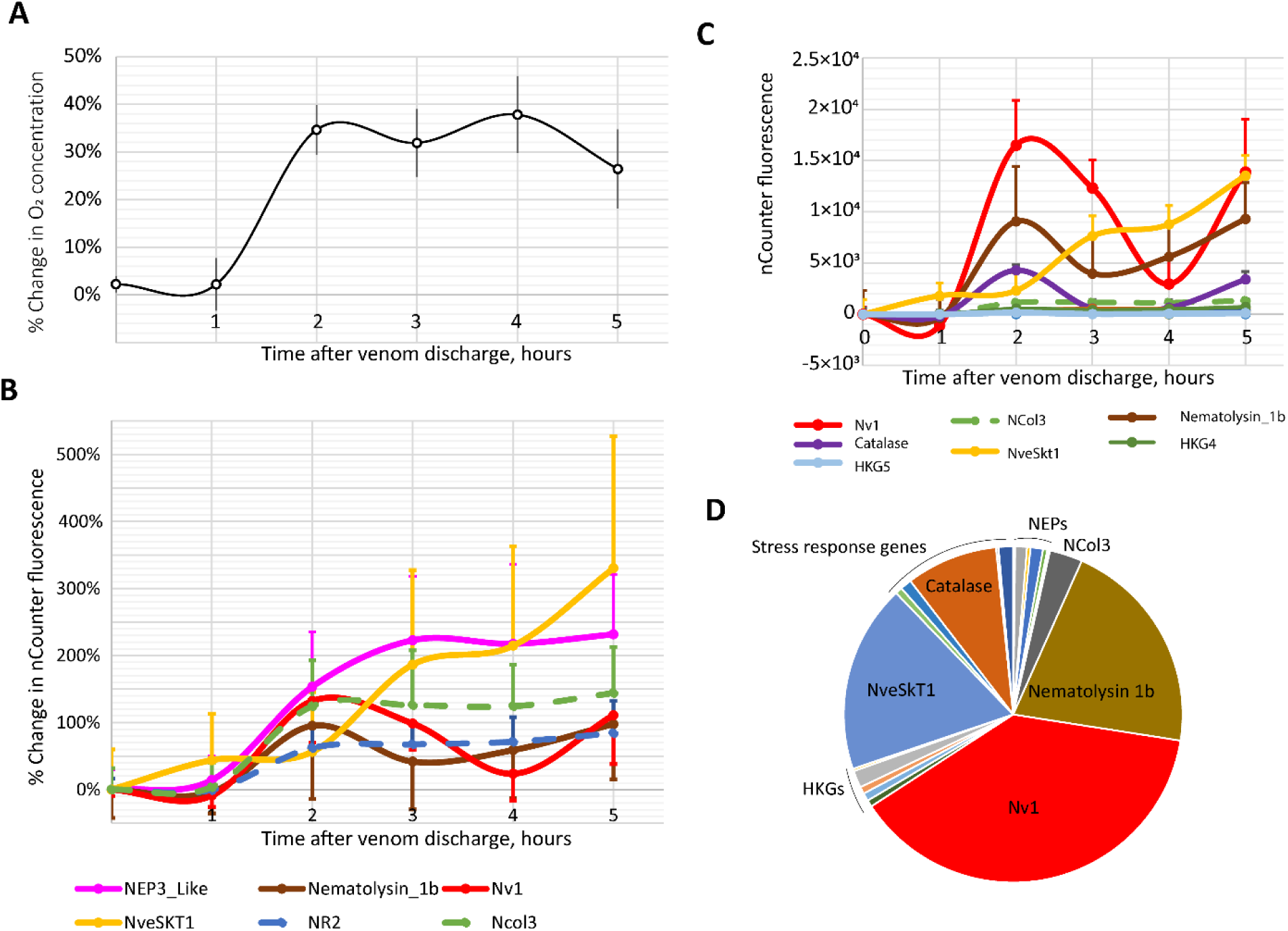
Venom production is metabolically expensive. **A)** Changes in oxygen consumption following fishing line treatment measured by closed-chamber respirometry. **B)** Percentage differences of expression for genes encoding toxins and nematocyst structural proteins following fishing line treatment. **C)** Absolute differences of expression for toxin-coding genes, nematocyst structural proteins and housekeeping genes following fishing line treatment. Noticeably, Nv1, NveSkT1b and Nematolysin 1b are the genes with the largest increase in expression in absolute units. **D)** Gene expression at the 3 h time point represented as a percentage chart. Nv1, NveSkT1b and Nematolysin 1b are contributing to a total of 77% of the transcripts among all the genes measured in this experiment. In **B, C**, and **D**, gene expression is represented as normalized fluorescence units measured by nCounter technology.

To confirm that the elevated metabolic rate involved the biosynthesis of venom, we repeated the venom depletion treatment and measured expression levels of genes encoding toxins produced by both adult males and females, nematocyst structural proteins, genes involved in general stress responses (e.g., cytoplasmic heat shock proteins and superoxide dismutases), and several housekeeping proteins by nCounter technology (**Suppl Data 1**). In addition to previously characterized venom components, we also included the putative toxins NEP8-like and NveSkT1 that have not been reported earlier (**Table 1, Suppl fig 1**). After two hours we observed an increase in the expression of all the genes measured (**Fig 1B**). At the 3h-5h time points, expression levels had diverse dynamics: several genes encoding toxins and nematocyst structural proteins (NveSkT1, NEP6, NEP3-like, NEP8, NR2, NCol3) kept increasing while others (Nv1, NEP14) showed wave-like fluctuations. Because the basal expression levels of the Nv1, Nematolysin1b and NveSkT1 toxins were comparatively high, this twofold increase in transcription contributed to the overall transcriptional activity much more than any other gene measured (they account for 77% of the transcripts among the genes measured at the 3 h time point) (**Fig 1 C, D**). Thus, increased expression of venom components and nematocyst structural genes positively correlates with the increase in respiration rate.

**Table 1.**
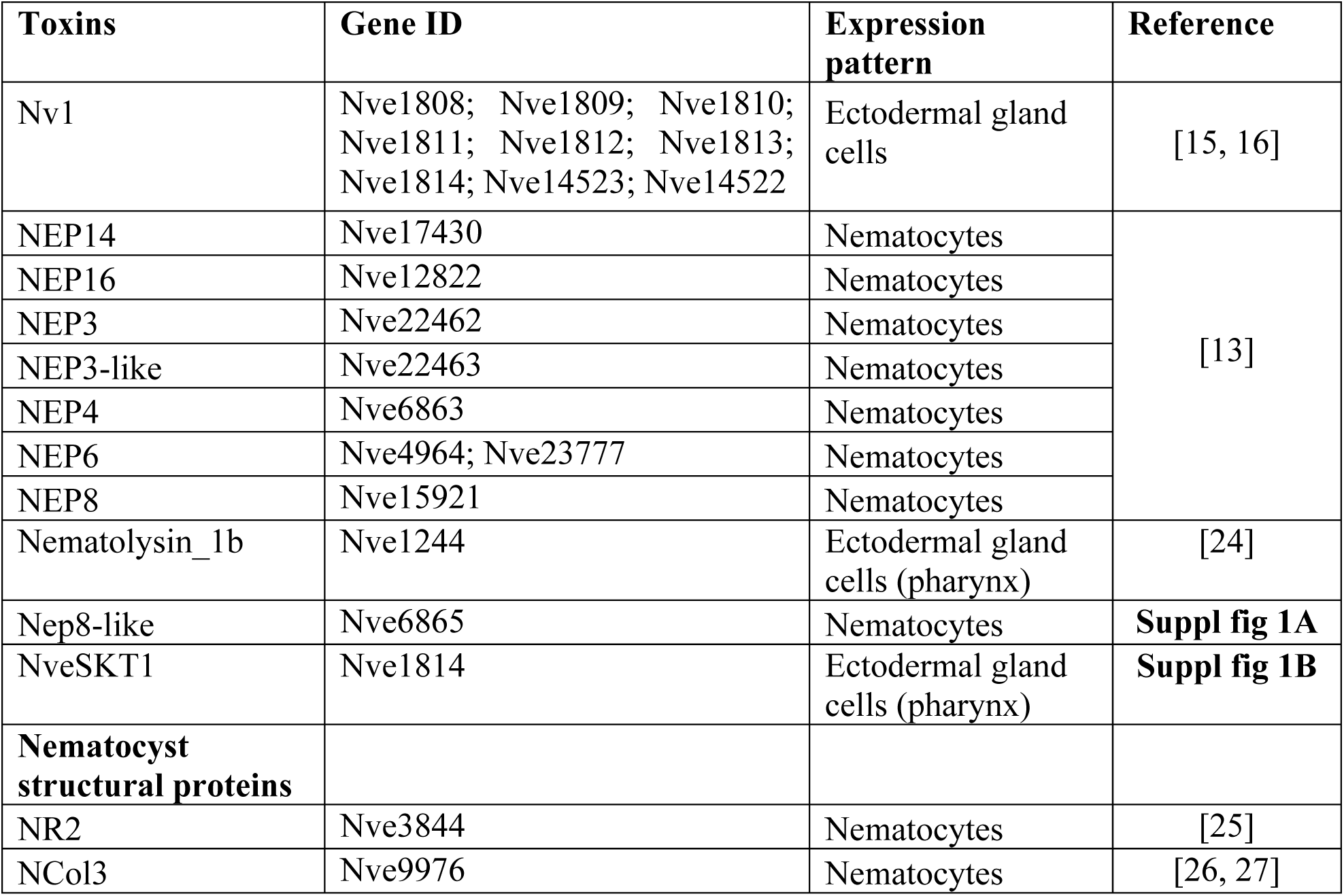
Venom components studied in the present work. Gene models are taken from https://figshare.com/articles/Nematostella_vectensis_transcriptome_and_gene_models_v2_0/807696

### Study of toxin production under stress conditions

*Nematostella* polyps were sampled from five locations distributed from the north to the south along the North American Atlantic coast (Nova Scotia (NS), Maine (ME), New Hampshire (NH), Massachusetts (MA), and North Carolina (NC); **Fig 2A, B**) and then cultured in the laboratory under common garden conditions. To determine the regulation of toxin production in response to abiotic stresses, we subjected adults from each location to the heat stress of 28 °C and 36 °C, UV light, low salinity (5‰) and high salinity (45‰; MA and NC only) water. Expression levels of toxin and stress response genes were measured by the nCounter platform [28] (**Suppl Data 2, 3**). To assess the biological significance of the expression changes in this experiment, we set a threshold of 2.4-fold change (corresponding to the maximum change in the expression of the normalizing housekeeping genes (HKG) [12]). In **Suppl tables 1** and **2**, changes in gene expression that are both biologically and statistically (p<0.05, Student’s T-test) significant are highlighted in bold red.

**Figure 2.**
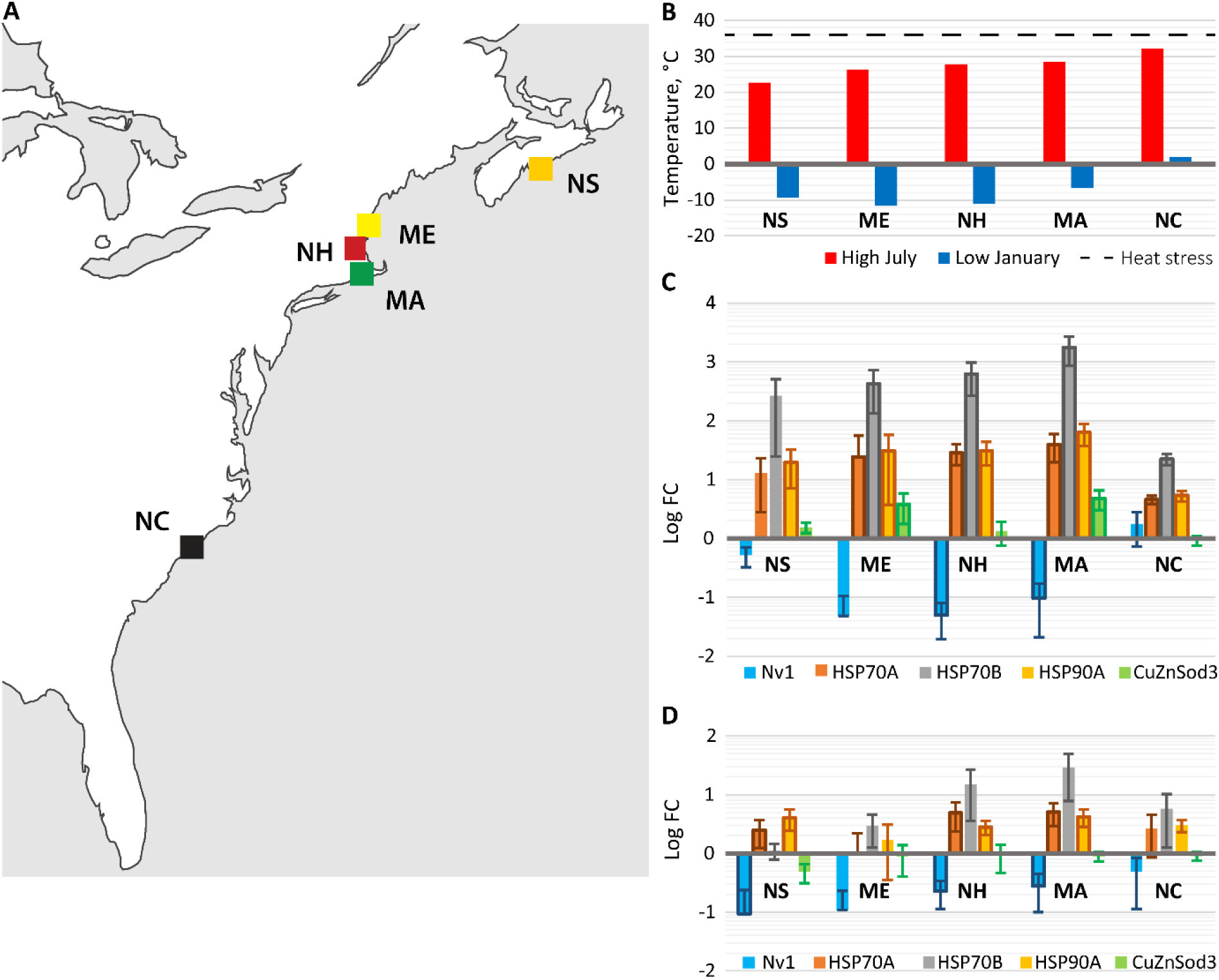
*Nematostella* populations from different climatic conditions respond differently to heat stress. **A)** Distribution of *Nematostella* populations along the East coast of North America. **B)** Average high July (red) and average low January (blue) temperatures across the populations from the North to the South. The data originate from publicly available sources (https://www.usclimatedata.com/climate, https://en.climate-data.org/north-america/) for the cities closest to the sampling points (Halifax, NS; Portland, ME; Manchester, NH; New Bedford, MA; Wilmington, NC). **C)** and **D)** Gene expression dynamics under heat stress (+36°C) **(C)** and low salinity stress (5 ‰) **(D)** among the populations from the North to the South. If the change in expression of a gene is greater than 2.4 times and p<0.05 (Student’s T-test), the corresponding data point is outlined in bold. Only genes with biologically significant expression change at least in two samples are shown.

The lower temperature of 28 °C, high salinity and UV light exposure did not result in a substantial change in toxin expression (**Suppl table 1, 2; Suppl fig 2**). However, under the high heat stress of 36 °C, we observed changes in the expression of toxins and HSPs. In the ME, NH, and MA populations, Nv1 production decreased beyond the threshold and up to 25-fold in the NH population compared to the control conditions (20 °C) (**Fig 2C**). In the ME population, the difference was not statistically significant (p>0.05, Student’s T-test). Additionally, NEP14, NEP16, NEP4, NEP8-like, and Nv1 toxin genes exhibited reduced expression under heat stress in the MA population (**Fig 3 A, B**). In the NS and NC population, expression level of Nv1 and NEP toxins did not change more than the threshold of 2.4-fold. HSPs expression showed interesting dynamics (**Fig 2**): the expression change was less pronounced in the most northern NS population (most of the changes were not statistically significant), then increased in ME, NH, and MA and then it dropped in NC. Thus, HSP expression showed an opposite trend to Nv1 expression. Physiological stress induced by low salinity resulted in decreased Nv1 expression in NS, NH and MA populations with the highest change in NS sample (10-fold) (**Fig 2D**); however, other toxin genes were not affected (apart of small change in NEP16 in MA). Expression of HSP70A and HSP90A increased in NS, NH and MA.

**Table 2.** Design of individual stress condition experiment. Numbers indicate the number of replicates for each condition making up a total of 87 samples.

|  | Control | 28 °C | 36 °C | 5‰ | UV light | 45‰ |
| --- | --- | --- | --- | --- | --- | --- |
| *nCounter run #1* |  |  |  |  |  |  |
| NS | 3 | 3 | 3 | 3 | 3 | 0 |
| ME | 3 | 3 | 3 | 3 | 3 | 0 |
| NH | 3 | 3 | 3 | 3 | 3 | 0 |
| MA | 3 | 3 | 3 | 3 | 3 | 0 |
| NC | 3 | 3 | 3 | 3 | 3 | 0 |
| *nCounter run #2* |  |  |  |  |  |  |
| MA | 3 | 0 | 0 | 0 | 0 | 3 |
| NC | 3 | 0 | 0 | 0 | 0 | 3 |

**Figure 3.**
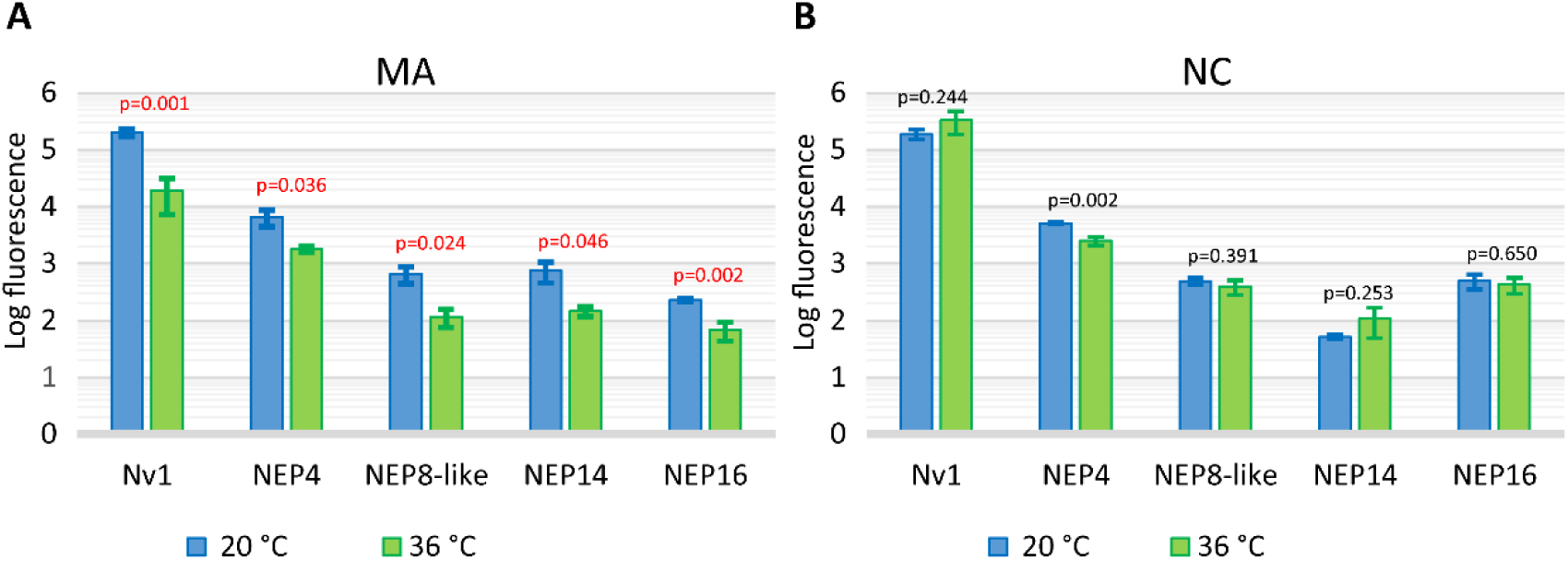
Expression of toxins in Massachusetts (A) and North Carolina (B) following heat stress. Gene expression is represented as Log10 of normalized fluorescence units measured by nCounter technology. P-values calculated by Student’s t-test are shown for each gene. If the change in expression of a gene is greater than 2.4 times and p<0.05, the corresponding p-value is shown in red.

To dissect regulation of venom production further, we focused on the MA and NC populations, which showed significantly different responses to temperature and had similar Nv1 expression levels under control conditions (**Suppl fig 3**). According to temperature data from publicly available sources, MA and NC also have quite different temperature profiles, including for example, the difference of 3.7 °C between high average July temperatures (**Fig 2B**). The different thermal regimes between MA and NC habitats are supported by our temperature monitoring data acquired by loggers which had been placed into ponds in March 2016 and measured temperature until December 2016 (**Fig 4A**). In the NC location temperature reached 36°C or higher on 110 days while in MA only on 38 days during the monitored period (**Fig 4B**).

**Figure 4.**
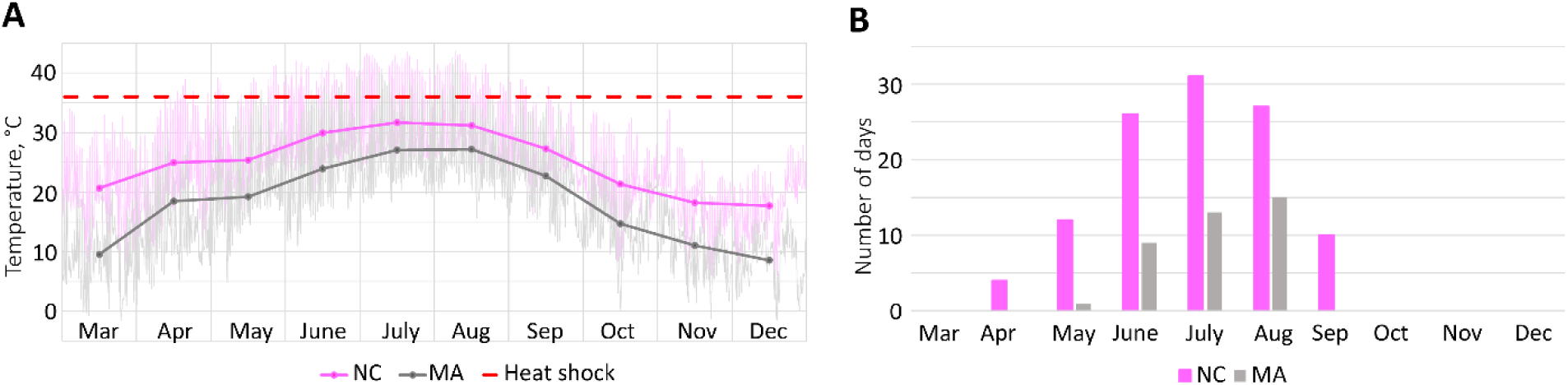
Thermal regimes differ between *Nematostella* habitats in Massachusetts and North Carolina. **A)** Water temperature in native *Nematostella* habitats recorded in March – December 2016. Measurements were taken every 20 minutes by a temperature logger. **B**) Number of days when the water temperature reached 36 °C or higher in March – December 2016.

### Transcriptome Annotation and Gene Ontology Analysis of the Stress Response

*Nematostella* in its natural environment, like most organisms, experiences stressors in combinations. We compared these former responses to temperature, salinity, or UV light with a combination treatment of all the three abiotic conditions. Adults from NC and MA were subjected to high temperature (36 °C, 24 h), UV light (6 h) and high salinity (40‰, 24 h). Control conditions were 20 °C, 15‰ and darkness. Animals were snap frozen (4 per sample), RNA was extracted and Illumina RNA-seq was performed in parallel for stress and control conditions.

Our differential expression analysis identified 3,637 transcripts that were differently expressed between controls and experimental groups for MA and NC populations (**Fig 5B**). Under the combined stress conditions, 738 genes were differentially expressed in both populations, 542 genes – in NC and 592 genes in MA (**Fig 5A, Suppl tables 3 – 5**). Within NC population *Nematostella* had 1,280 transcripts that were differentially expressed, 457 of which contained Gene Ontology (GO) information comprising of 4,667 different GO groups across these transcripts. For the MA population there were 1,330 transcripts that were differentially expressed, 513 of which contained some GO information comprising of 5,405 different GO groups across these transcripts. The largest proportion of the differentially expressed GO groups for NC animals were associated with the Biological Process domain (79.7%), while Molecular Function (15.4%) and Cellular Component (8.8%) domain were associated with proportionately fewer transcripts. Similarly to the NC population, in the MA samples the largest proportion of the GO groups were associated with the Biological Process (81.1%) GO domains, while Molecular Function (12.1%) and Cellular Component (6.8%) were associated with proportionately fewer transcripts.

**Figure 5.**
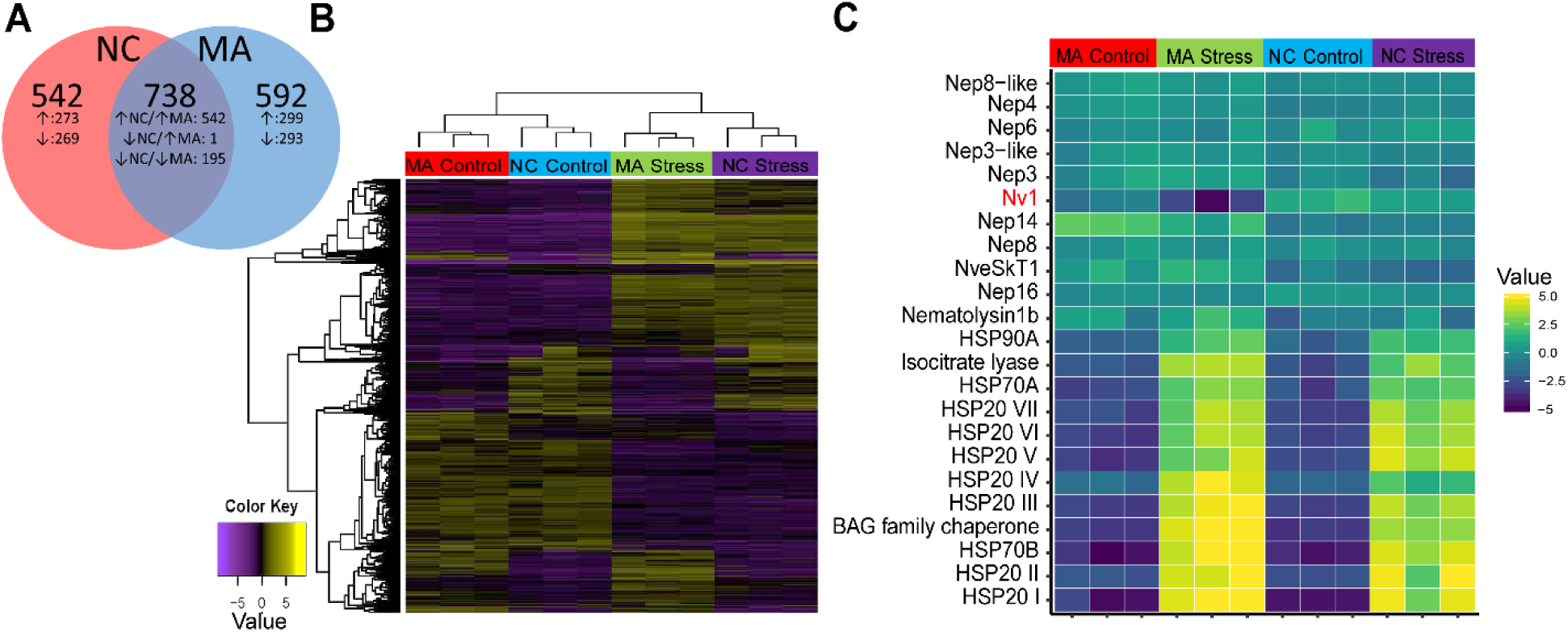
Differential gene expression among Massachusetts and North Carolina populations after “combined” stress (+36 °C, UV light, water salinity of 40 ppt). **A)** Venn diagram of differentially expressed genes that are shared and unique for the MA and NC populations. **B)** Heatmap of differentially expressed genes across treatments and populations. **C)** Heatmap of toxin genes and stress response genes. The latter are the most differentially expressed genes between the control and stressed conditions in both populations.

We used comparative GO analysis to identify functionally important GO groups in relation to environmental stress response. However, our analysis did not identify any characterized functionally important groups more highly represented than other GO groups (**Suppl Table 6**). The GSEA analysis was able to more accurately identify GO groups that were variable across treatments. In both populations, the “response to heat” GO term (GO:0009408) was enriched (**Suppl Table 7**). The most differentially expressed genes between the control and stressed conditions for both populations can be traced back to stress response genes such as HSP20 and HSP70 (**Figure 5C, Suppl Table 8**). Beyond these stress response genes, our GSEA also revealed an enrichment in the upregulation of transcription factors (TF) in both populations following stress response. Differences in upregulation of specific transcription factors were observed among the populations, with 30 TFs upregulated in both populations, 27 TFs upregulated only in MA, and six TFs upregulated only in NC **(Suppl Table 9)**.

In the control groups, many of the differentially expressed transcripts had relatively low levels (TMM<10), therefore, even a small change in gene expression resulted in a dramatic fold change. When considering LogCPM (Log counts per million), which accounts for more highly expressed transcripts overall, strikingly, the highly expressed Nv1 toxin gene (TMM=847±243 in MA; TMM=3824±1112 in NC) is the most downregulated locus in the MA population.

### Northern (MA) but not southern (NC) Nv1 expression reacts to combined stress conditions

Among toxin genes, only Nv1 in MA showed a substantial change under stress conditions, where it decreased 5.5 times, p = 0.01 (**Suppl Table 8, Fig 5C**). Expression of NveSkT1, Nematolysin1b, nematocyst toxins (NEP3, NEP4, NEP3-like, NEP8, NEP14, NEP16, NEP6) and structural proteins (NCol3 and NR2) remained more stable in both populations as changes were in the range of 0.4 – 2.1-fold, and in most cases were not statistically significant. Among the stress response genes that we analyzed under individual stress conditions, heat shock protein genes (HSP70B, HSP70A, HSP90A) exhibited an increased expression level of 11 to 821 fold. In contrast, the expression of the CuZnSod3 superoxide dismutase gene remained stable while MnSod1 decreased approximately twofold (**Fig 5C, Suppl Table 8**).

To study the effects of the stress conditions on Nv1 production at the protein level we used LC-MS/MS (**Fig 6, Suppl Data 4**). It was noticeable that control LFQ values decreased between our measurements apparently due to degradation or aggregation of Nv1 peptides. Thus, to control for this technical variation, the samples were run in pairs of control and stress for each population in triplicates. In each pair from MA, the LFQ value for the stress sample was approximately twice lower than in the control. On the other hand, in NC, stress LFQ values were slightly higher than control. Thus, the observed trend supports our findings on the transcriptomics level.

**Figure 6.**
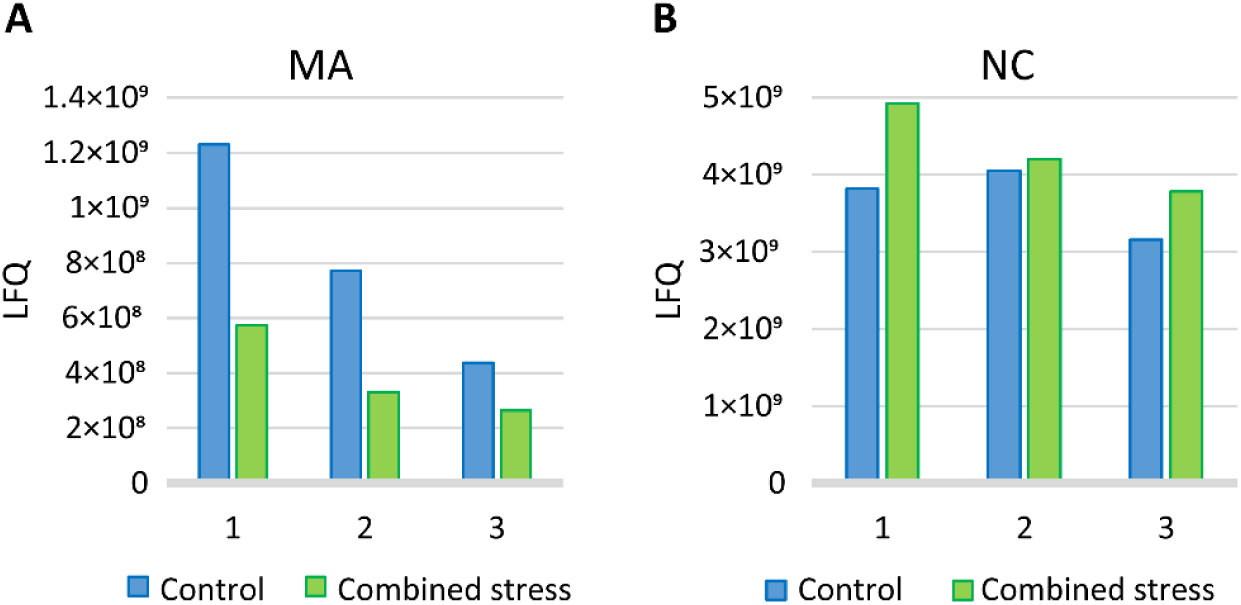
Nv1 production at the protein level under control and “combined” stress conditions measured by LC-MS/MS in Massachusetts (A) and North Carolina (B) populations. The numbers correspond to technical replicates.

## Discussion

In the current study we provide a suite of molecular and physiological evidence that toxins are metabolically costly to produce for a venomous species and that individuals from different populations along a thermal gradient exhibit unique modulations in toxin expression when experiencing a range of abiotic stressors.

One of the significant findings for this research is the first substantial evidence for a cost of venom production in a cnidarian. We have identified that an increase in expression of venom components and nematocyst structural genes correlates with increase in respiration rate demonstrating that venom biosynthesis is metabolically expensive in *Nematostella*. Toxin proteins have been hypothesized to be energetically expensive to produce and thus venomous species should be under strong selection to modulate the quantity and composition of venom cocktails [18, 29]. Studies in reptiles and arachnids [18] support this hypothesis where individuals modulate feeding preference or behavior depending on their energetic state or the relative venom quantities (where starved individuals produce less venom). This hypothesized tradeoff has some previous support by two anecdotal observations in cnidarians: stony corals strongly downregulate under heat and UV stress the expression of their small cysteine rich peptides (SCRiPs) [30], which we later found to be neurotoxins [31], and among the genes that the sea anemone *Anthopleura elegantissima* downregulates under similar stress conditions were genes encoding the Nv1 homologs Anthopleurin-C (called “Toxin PCR 7” in that study) and APETx1 [32]. Thus, adjusting venom production to the metabolic status may be an evolutionary advantage in cnidarians in general. Our data from *Nematostella* indeed show that among venom components, the highly abundant toxin Nv1 is the most downregulated under heat stress conditions in multiple populations. The investment of energy into heat response (e.g., production of numerous heat shock proteins, HSPs) appears to be traded-off with other high cost physiological processes that do not contribute to survival under heat stress (e.g., venom production).

In sea anemones venom release involves secretion of venom components from ectodermal gland cells [15] and discharge of single-use nematocysts. In our experiment, after the mechanical stimulation venom components need to be replenished and also new nematocytes have to maturate and produce nematocysts. Therefore, multiple processes involved in venom regeneration may explain the response where venom-related genes as well as other genes were upregulated after two hours following the treatment.

Another significant finding in the current study is the result that *Nematostella* polyps from different geographic locations differ in their production of toxins when experiencing the same environmental stressors. While we measured significant changes in expression of multiple toxins under different exposures to high temperature, low and high salinity, and UV light, the largest population-specific impact was at the higher temperature exposure. Responses to the heat shock followed an inverted bell-shaped profile along a latitudinal temperature gradient: decrease in Nv1 expression was observed in the NH and MA populations but not in the NS and NC populations (**Fig 4**). One of the possible explanations would be adaptation to the local temperature conditions for these locations. In NC, where the average high July temperature is 32.1 °C and average low January temperature does not go below 0 °C, *Nematostella* would be exposed to heat shock of 36 °C regularly. On the other hand, in NS the average high temperature is 22.6 °C meaning that *Nematostella* is very unlikely to experience a heat shock of 36 °C in this habitat and would be rather adapted to a lower temperature ranges. The lack of a gene expression response in NS animals may be due to lack of adaptations to such an extreme temperature compared with typical high temperatures causing severe stress and a general metabolic shutdown. In ME, NH and MA the average high temperatures are between 26.2 to 28.4 °C and low temperatures are below 0°C indicating that *Nematostella* would be exposed to both high and low temperatures and thus is adapted to more diverse temperature regime. Our temperature monitoring data in *Nematostella* habitats confirm that NC population may be exposed to 36 °C for six months each year while the MA population only during four months. We hypothesize that heat response mechanisms in all populations are inducible based on the expression of HSP genes, but there is only a tradeoff for physiological processes in NH and MA populations. The NC anemones have potentially evolved other physiological mechanisms to cope with higher temperatures and to avoid the tradeoff with the production of toxin proteins. However, it is possible that for this population a tradeoff might be occurring at even higher temperatures. Similar mechanism of plasticity in environmental stress response was described in the coral *Porites astreoides* [33].

Interestingly, the combined stress conditions (heat, UV light, and high salinity) provoked a similar response as heat stress alone: Nv1 was downregulated in MA but not in NC. This result indicates that single stressors (temperature) and combined stressors induce a similar transcriptional response in the most abundant venom component, Nv1. Analysis of the changes in functional Gene Ontology groups under stress between MA and NC populations revealed that, in concordance with the previous lines of evidence, heat shock proteins underwent significant upregulation in both populations. Differences in the regulation of Nv1 biosynthesis may be explained by a number of transcription factors differentially upregulated between the populations under stress. Factors underlying adaptation of NC population to retain high venom production levels under stressful conditions are yet to be discovered. Despite the fact that multiple genes were differentially expressed between the populations, our analysis of enriched GO terms did not reveal any specific functional pathways upregulated in NC population and potentially keeping venom production beneficial or neutral under the stress. One of the reasons might be incomplete annotation of cnidarian genomes and GO terms due to high divergence from bilaterian organisms with well-annotated GO terms such as vertebrates, *Drosophila melanogaster* and *Caenorhabditis elegans*.

Our earlier and present work have demonstrated that Nv1 toxin is regulated by diverse environmental and endogenous factors: abiotic stress, light/dark cycles and life stages. Unlike other venom components, it is encoded by multiple gene copies potentially underlying the high expression level and providing higher regulatory flexibility. Because responses to the abiotic stress differ between populations experiencing a gradient of environmental conditions, our data suggest that regulation of venom expression is adjusted and potentially “optimized” to conditions reflective of their local environments. Thus, modulation of venom production appears to be one of the evolutionary adaptations in *Nematostella*. Adaptation of this trait is plausible as we showed that production of venom and venom-delivery cells is metabolically expensive and the balance between the benefits and costs of venom production might change dramatically between habitats due to different abiotic conditions.

## Materials and methods

### Venom discharge and oxygen consumption

To provoke venom discharge, *Nematostella* polyps were subjected to mechanical stimulation with a gelatin-coated fishing line imitating interaction with predators and prey [34]. Closed-chamber respirometry is an effective means to measure oxygen consumption to determine shifts in metabolic rate for animals in different conditions. We used comparative respirometry to measure changes in oxygen consumption in anemones following mechanical stimulation with a fishing line. Anemones were placed in 3 ml water-jacketed respiration glass chamber, given approximately 30 minutes to open their tentacle crown, and then the tentacles were probed with the monofilament line. The respiration chambers were then closed and measured continuously for five hours. Oxygen uptake by individual anemones was measured using Clarke-type oxygen electrodes (YSI, USA). Two-point calibration of electrodes was performed before each day, and continuous data acquisition of oxygen concentrations was made using a BIOPAC Data acquisition system (BIOPAC, USA). Time effects on respiration were determined during the time course and compared with anemones that received no physical stimulus.

To confirm that respiration changes correlated with changes in expression of venom genes, venom discharge treatment was repeated and animals were sampled and frozen in liquid nitrogen every hour for 5 h for RNA extraction. Every time point was sampled in triplicates, 3 animals/replicate.

### Stress conditions

Before treatments, animals from NS, ME, NH, MA and NC populations were acclimatized at 20 °C for 24 h in the dark in 15‰ artificial sea water (ASW). For the individual stress conditions, three groups of animals were exposed to 28 °C, 36 °C, 5‰ ASW, UV light, 45‰ ASW (only MA and NC), or control conditions (20° C, 15‰, dark) separately. Animals were placed into tubes and frozen to obtain 3 replicates for each condition, 2 animals/replicate, total 87 samples (**Table 2**). For the combined stress treatment, half of the acclimatized animals from MA and NC populations were transferred into 40‰ ASW and exposed to 36 °C for 24 h. For the first 6 h of treatment, the UV lamp was on. Control animals were kept at 20 °C for 24 h in dark in 15‰ ASW. After the treatment animals were placed into plastic tubes and frozen in liquid nitrogen to make three replicates for each condition and 4 animals per replicate, total 12 samples.

### Quantification of gene expression

Total RNA was extracted from the frozen samples using RNeasy Mini Kit (Qiagen, Germany). RNA quality was assayed with a Bioanalyzer NanoChip (Agilent Technologies, USA).

For the samples collected following venom discharge and individual stress samples, the nCounter platform (NanoString Technologies, USA; performed by Agentek Ltd., Israel and MOgene, USA) was used. 100 bp probes (**Suppl Table 10**) were designed to specifically bind to transcripts encoding toxins, nematocyst structural proteins and stress response genes. For normalization of expression, a geometric mean of expression levels of five genes with stable expression across development was used (similarly to [12, 14]). Fold change and absolute change relatively to the 0 h time point were calculated for the samples collected at 1 h, 2 h, 3 h, 4 h, 5 h time points after venom discharge. Additionally, at the 3 h time point expression level relatively to the total expression of all the genes measured in this experiment (100%) was calculated and represented as a pie chart (**Fig 1 D**). For the 87 individual stress samples, the measurement was performed in two batches as indicated in **Table 2**. For each population (for each nCounter batch separately), every stress condition was compared to the control and p-values (Student’s T-test) and fold changes were calculated.

For the 12 “combined stress” samples, cDNA libraries were constructed by the Sense kit (Lexogen, Austria), pooled together and sequenced by the NextSeq500 platform (Illumina, USA) with 400 million read depth and 40 bp paired end reads (performed at the Center for Genomic Technologies, The Hebrew University of Jerusalem). Raw reads were submitted to the NCBI Sequence Read Archive database (Submission ID: SUB6850129, BioProject ID: PRJNA601530). The raw reads were mapped to *Nematostella* gene models [35] by Star aligner [36]. Differential expression analysis was performed by EdgeR Bioconductor package [37]. This allowed us to identify transcripts with significant deviations in expression level. In our EdgeR analysis, we identified transcripts that were differentially expressed across all treatments in combination, as well as within each focal locality. Both metrics of fold change and LogCPM were used to identify any potentially informative transcripts for further evaluation.

### Proteomics

For measurements of toxin production at the proteomics level, only females from MA and NC populations were used. After the “combined stress” treatment, the animals were frozen (3/tube) and lysed in 8M urea, 400 mM ammonium bicarbonate. The lysates were centrifuged (22,000 × g, 20 min, 4°C), protein concentration was measured with BCA Protein Assay Kit (Thermo Fisher Scientific) and 10 µg aliquots of protein were sent for LC-MS/MS analysis by Q Exactive Plus mass spectrometer (Thermo Fisher Scientific) at the Proteomics Center of the Alexander Silberman Life Sciences Institute, The Hebrew University of Jerusalem. All the procedures were performed as described by [12]. The samples were analyzed in technical triplicates. The mass spectrometry proteomics data have been deposited to the ProteomeXchange Consortium [38] via the PRIDE [39] partner repository with the dataset identifier PXD016943.

### Transcriptome Annotation and GO analysis

Using information derived from the previously predicted and annotated gene models [35] we evaluated whether differentially expressed transcripts may have carried both functional information based on TMM normalized fold change values as well as LogCPM values. TMM normalization aides when scaling for variation in library size as well as transcript diversity being sampled [40]. The use of TMM normalization fold change alone aided in identifying transcripts expressed at very low levels in the control groups in order to identify potentially functionally important transcripts. In contrast, LogCPM is also a measure of log counts per million contrasting control and treatment, providing more weight to differences in gene expression when they’re more highly expressed across all treatments.

Beyond individual transcripts we evaluated changes in functional Gene Ontology groups (GO) between the different populations across control and stressed environments. This was done by two different approaches, first we used custom Python scripts as previously described in [41], which divided TMM values across associated GO terms. The GO groups were then grouped based on semantic similarity and identified using REVIGO [42]. The second approach we performed was gene-set enrichment analysis (GSEA) on the differentially expressed genes using GOseq [43]. This required genes to first be annotated against the Swiss-Prot database (accessed 27/10/19) using BLASTp and gene ontologies mapped.

## Acknowledgements

We would like to acknowledge Dr. Anna Ivanina (UNC Charlotte) for assistance with the design and completion of oxygen consumption experiments. We would also like to acknowledge the following researchers of the core facilities of the Alexander Silberman Institute of Life Sciences, The Hebrew University: Dr. William Breuer (Interdepartmental Unit) for his help with mass spectrometry and Dr. Michal Bronstein and Ms. Adi Turjeman (The Center for Genomic Technologies) for their help with high-throughput sequencing.

