## Supplementary figures for "Population Specific Adaptations in Venom Production to Abiotic Stressors in a Widely Distributed Cnidarian"

### Supplementary figure 1. New venom components

**1A. NEP8-like is expressed in nematocytes in larvae and primary polyps and shares high sequence similarity with NEP8 toxin found in nematocysts.** Signal peptide is shown in bold, ShKT domains are highlighted, and custein residues forming the ShKt motives are in green.

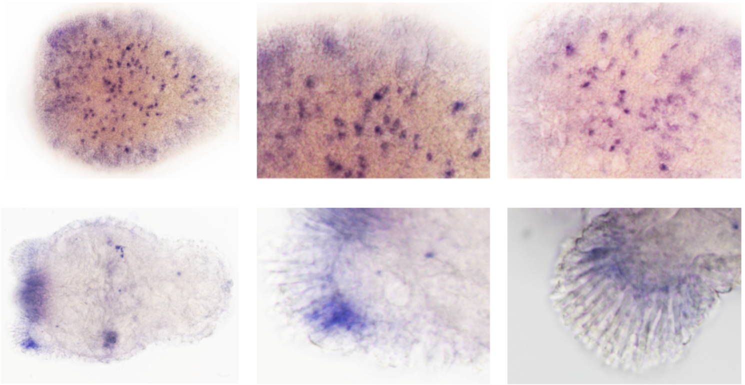

```
Nep8_Nve15921           MLRRPLLLVLFTVFSTLYAKDLRGVSPPTNESEAEVSPGDDEGPPEPGNEPDVNWRTLTI 60
Nep8_like_Nve6865       MASLFWLLVACLVLLAVDAKEIRRKKLYRQ--------------HESLRTIAETLERVHI 46
                        *     ***   *: :: **::*  .   :               *  .    . . : *

Nep8_Nve15921           DPEDCKDKGKDCESLADEGNCLKKLNYAVGNCPWTCRFCKKEN-GDSKKCKDLAGERHCN 119
Nep8_like_Nve6865       SPEDCKDYKSNCEEFADEGSCLKRFDYGFKFCPWSCRFCAKPEKYNTERCADLAGK-RCG 105
                        .******  .:**.:****.***:::*..  ***:**** * :  ::::* ****: :*.

Nep8_Nve15921           GWKVKGDCVRLPDYMMQNCKLSCELCGPETRFKYTDEDVRCPEWAQAGYCSTNTDINLKC 179
Nep8_like_Nve6865       PWAQRSECLRLPEFMQQNCKQSCDLCGPENA--YTDDDVRCPYWGKEGYCRTDAEINSRC 163
                         *  :.:*:***::* **** **:*****.   ***:***** *.: *** *:::** :*

Nep8_Nve15921           PHSCRKYKQRAPASTPYPYPVEALHPYQYRVVLPSVTILQTTTA-----APSTQPAETT- 233
Nep8_like_Nve6865       PYSCRKYRSVSPAPQPYPYPIGTLYPYQPTPVP-GVTVSQPKIVLQPYPAPYPYPYPVPS 222
                        *:*****:. :**  *****: :*:***   *  .**: * . .     **   *  .

Nep8_Nve15921           KAPPNTAAPTAAP------------TPAPTPAPAPAPTPAPAPAPTPAPAPAPATTPAPA 281
Nep8_like_Nve6865       PAPSPSPSPSPAPSPSPSPSPALSPSPAPSPSPSPSPSPSPSPSPSPSPSPSPSPSPSPS 282
                         **  : :*: **            :***:*:*:*:*:*:*:*:*:*:*:*:*: :*:*:

Nep8_Nve15921           PVP---PAPAPAPAPAPAPP-APPVAPAPQTAPLAGSPPESTPEEQDD--------NSA- 328
Nep8_like_Nve6865       PSPEPSPSPAPSPAPEPSPAPSPASEPSPETTTPAPKDPTQAPVVVETQQPSTQLPAPTQ 342
                        * *   *:***:*** *:*  :*   *:*:*:  *  . * .:*   :           :

Nep8_Nve15921           DESTEI----EAGEGGGELCDEKHSSQQ-------------------------------- 352
Nep8_like_Nve6865       PPTEPIDSSGNPGESGNGVWPGQKGQPSAAPVTQPQGSGEEEEVEEGVTTDAPVGSPVPE 402
                          :  *    : **.*. :   ::.. .

Nep8_Nve15921           ------------------------------------------------------------ 352
Nep8_like_Nve6865       ATPGPAATNAPEATQAPEVVTQAPEATSAPEVTAAPTGQATPASNPIGTPPVVFDHRKSH 462

Nep8_Nve15921           ------------------------------------------------------------ 352
Nep8_like_Nve6865       RSRNAKNDAHHHNNNNHHNNNNHHNNPHNNAPSTPPTAPPSPAPVFSKKLPVSGLDKATN 522

Nep8_Nve15921           -----------------  352
Nep8_like_Nve6865       AAKAKRNKHGHRHIISE  539
```

**1B. NveSkT1 is expressed in pharyngeal gland cells in planulae and primary polyps.** Signal peptide is in bold (identified by SignalP online tool, http://www.cbs.dtu.dk/services/SignalP/), peptydoglycan binding domain is in blue, metallopeptidase is in grey, and ShKT domain is underligned with cysteins in green (domains were identified by Interpro, https://www.ebi.ac.uk/interpro/). The ShKT domain od NveSkT1 shares noticeable sequence similarity with potassium channel blocker ShK from the sea anemone *Stichodactila heliantus*.

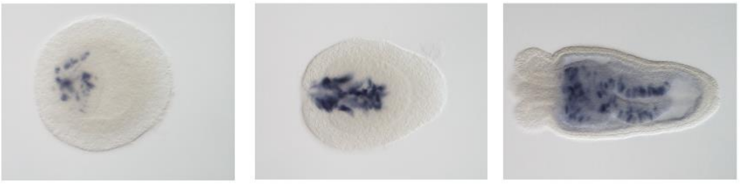

>NveSkT1

**MKTLVFAGLLCLIAFAVS**EDDDQTMALKYLNQFHYISPARSGNHNVKTALEKFQSFAGLPVTGEIDAATIA
QMKMPRCGMPDVDDDGLRIRRYKLGSKWNKKHLTYHISHGQDLSSSVQDRVFAKALDYWARVSGLTFSRTM
DGENADLKISFGPKSHGGTHDPEGTCSYPFDGPGGVLAHAFFPRNGRAHFDEDEDFTDGTYEGTNLLWVAT
HEFGHSLGLHHSDVRDAVMYPYYTGYKPGFDLKADDIAGIRAHYGDENSGGSQPCVDNNQSCPDWTIYCTN
NEYVIANCRKSCGSC

```
ShK_Stichodactyla        RSCIDTIPKSRCTAFQCKHSMKYRLSFCRKTCGTC   35
NveSkT1-ShKt_domain      -PCVDNNQSCPDWTIYC-TNNEYVIANCRKSCGSC   33
                           *:*.   ..    :: *   . :* :: ***:**:*
```

**Supplementary figure 2. Gene expression dynamics under UV light stress.**

**2A) Gene expression dynamics under UV light stress among the populations from the North to the South.** If the change in expression of a gene is greater than 2.4 times and p<0.05 (Student's t-test), the corresponding data point is outlined in bold.

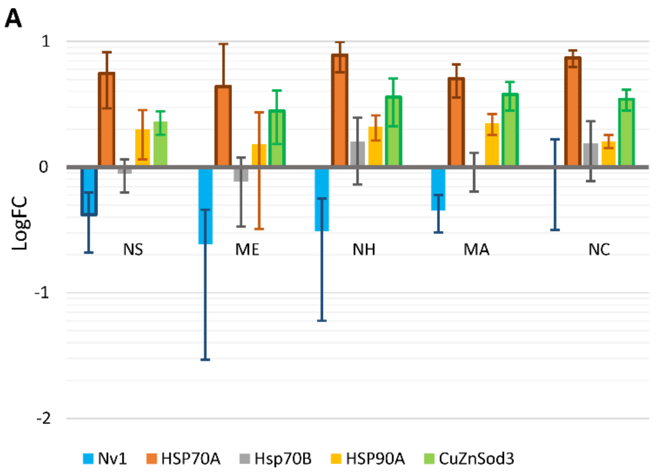

**2B and C) Expression of toxins in Massachusetts (B) and North Carolina (C) following UV light stress.** Gene expression is represented as $Log_{10}$ of normalized fluorescence units measured by nCounter technology. P-values calculated by Student's t-test are shown for each gene. If the change in expression of a gene is greater than 2.4 times and $p<0.05$, the corresponding p-value is shown in red.

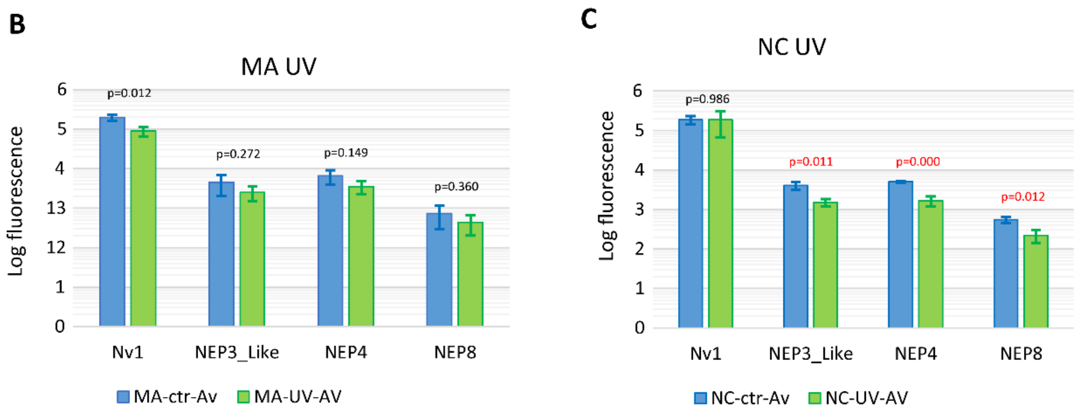


**Supplementary figure 3. Expression levels of Nv1 under control conditions in NS, NH, ME, MA, NC populations**

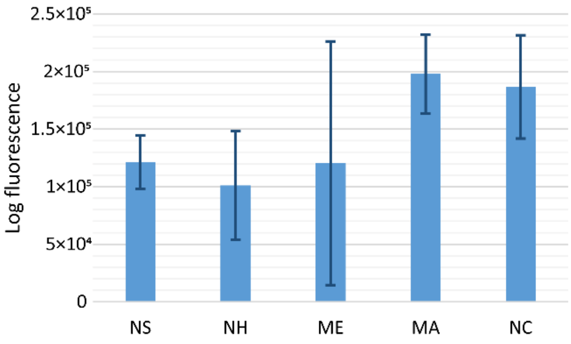
